# DPCGS: a computational framework for linking GWAS to single-cell transcriptomics in complex traits and diseases

**DOI:** 10.64898/2026.07.14.738331

**Authors:** Chonghui Liu, Bo Yuan, Baihan Shen, Jinjun Li, Ran Zhu, Pengpeng Yang, Bohan Wu, Yongquan Xuan, Siwei Yang, Naixue Yang, Longxuan Ma, Qiaoming Liu, Shaoxing Dai, Yan Zhang

## Abstract

Complex traits and diseases arise from the interplay between genetic variation and cellular heterogeneity, making it essential to understand how genetic risk manifests at the cellular level. However, connecting genome-wide association studies (GWAS) to specific cell populations remains challenging due to cellular complexity and the prevalence of noncoding variants. Here, we present DPCGS, a computational framework that systematically integrates GWAS summary statistics with single-cell RNA-sequencing (scRNA-seq) data to identify trait-associated cell subpopulations, genes, and regulatory programs. Unlike existing approaches that primarily evaluate pathway enrichment or cell-type-level associations, DPCGS quantifies the enrichment of genetically prioritized genes within individual cells through a statistically calibrated gene-set scoring strategy, enabling high-resolution mapping of genetic risk to cellular states. Benchmarking across simulated and diverse human single-cell datasets demonstrates that DPCGS achieves superior accuracy, sensitivity, and robustness compared with existing methods, including scDRS and scPagwas. Applying DPCGS to Alzheimer’s disease and asthma reveals disease-associated cellular populations and uncovers potential molecular drivers, including CD74, FOS, and AP-1 family regulatory programs, providing insights into disease-specific immune and cellular mechanisms. By bridging genetic discoveries from GWAS with functional interpretation at single-cell resolution, DPCGS establishes a generalizable framework for dissecting the cellular architecture of complex traits and diseases. This approach enables systematic discovery of disease-relevant cell subpopulations, regulatory networks, and potential therapeutic targets, offering broad applications in human genetics, single-cell biology, and precision medicine.

## Introduction

Complex traits and diseases such as Alzheimer’s disease (AD) and asthma are influenced by the interplay of genetic and cellular factors^1^. Genome-wide association studies (GWAS) have identified thousands of genetic variants associated with diverse traits, yet most of these variants lie in noncoding regions, making it challenging to interpret their functional roles^2^. A growing body of evidence indicates that genetic associations for a trait are often positively correlated with the expression of trait-relevant genes in bulk tissues or specific cell types^3, 4, 5^. This suggests that integrating genetic association data with transcriptomic profiles can help uncover the cellular contexts in which genetic risk manifests.

Single-cell RNA sequencing (scRNA-seq) has revolutionized our ability to study cellular heterogeneity by enabling transcriptomic profiling at single-cell resolution^6^. By dissecting the diversity of cell states within tissues, scRNA-seq provides a unique opportunity to link genetic risk factors from GWAS to disease-relevant cellular subpopulations. Identifying cell populations relevant to specific traits (including diseases) from scRNA-seq data is essential for elucidating the underlying mechanisms of complex traits and diseases^7^. Existing computational approaches, such as scDRS and scPagwas, provide initial frameworks for integrating GWAS signals with single-cell expression profiles, but they suffer from limited sensitivity, specificity, or scalability^8, 9^. As a result, the systematic identification of trait-associated cell subpopulations and their molecular drivers remains incomplete.

To overcome the limitations of existing approaches, we developed DPCGS (Identification of <u>D</u>isease <u>P</u>henotype-associated <u>C</u>ell subpopulations by integrating <u>G</u>WAS summary statistics with <u>S</u>cRNA-seq data), a computational framework that integrates genetic associations with single-cell transcriptomic profiles to identify trait-relevant cell populations, genes, and transcriptional regulators. Building on the principle that cells associated with a given trait exhibit stronger aggregate expression of trait-relevant genes than random gene sets, DPCGS prioritizes candidate genes from GWAS using the Multi-marker Analysis of GenoMic Annotation (MAGMA), quantifies their expression across individual cells to compute trait relevance scores (TRSs), and assesses statistical significance using matched control distributions. We systematically evaluated DPCGS on both simulated and real scRNA-seq datasets, demonstrating superior performance over existing methods in traits such as monocyte count and NK cell proportion. Furthermore, DPCGS identified disease-relevant cell subpopulations (oligodendrocytes and astrocytes in AD; macrophages and B cells in asthma) and potential regulatory genes such as *CD74, FOS*, and *FLI1*. Together, these findings establish DPCGS as a versatile framework for linking GWAS signals to single-cell expression variation, thereby providing new insights into the cellular and molecular mechanisms of complex traits and diseases and offering avenues for biomarker discovery and therapeutic development.

## Results

### Overview of DPCGS

Previous studies have established that genetic associations for a given trait are often positively correlated with the expression of trait-relevant genes in bulk tissues or specific cell types^10, 11, 12^. Building on this principle, DPCGS extends such analyses to the single-cell level by integrating scRNA-seq profiles with GWAS summary statistics. In brief, the method evaluates whether genes implicated by GWAS are expressed more strongly in certain cells compared with random gene sets.

To implement this, DPCGS first identifies putative trait-associated genes from GWAS data using MAGMA, which maps trait-associated SNPs to their proximal genes and prioritizes the top candidates (Fig. 1, top). It then calculates aggregate expression levels of these candidate genes for each individual cell to derive raw disease scores (Fig. 1, middle). To assess statistical significance, the method constructs matched control gene sets and generates distributions of control scores. The empirical p-value for each cell is computed using the distribution of pooled control scores derived from all control gene sets. (Fig. 1, bottom).

**Fig. 1.**
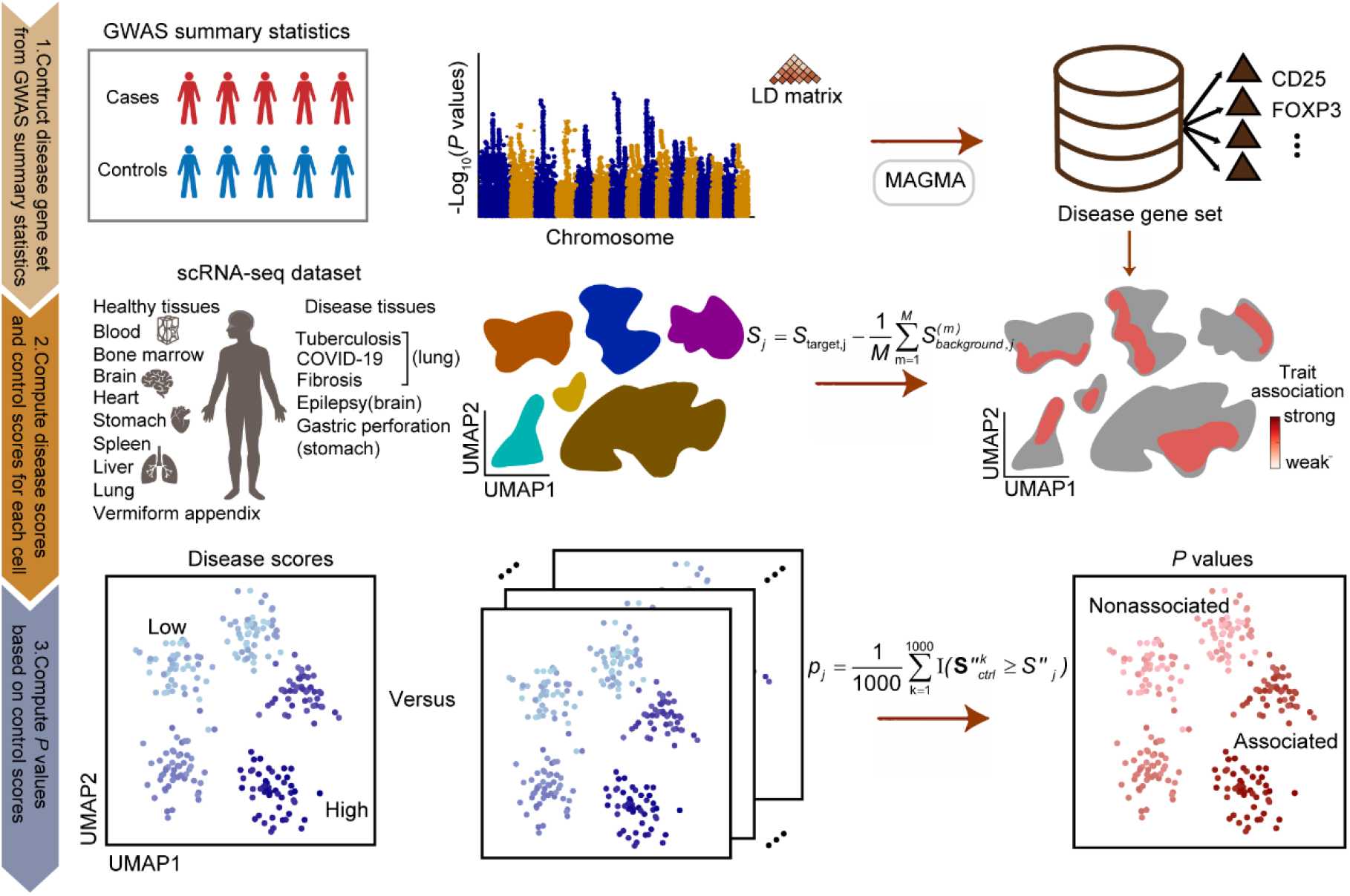
The workflow of the DPCGS. DPCGS integrates GWAS summary statistics with single-cell RNA-seq data to evaluate disease relevance at the cell level. Putative disease genes are first prioritized from GWAS summary statistics using MAGMA, with the top 1,000 genes selected as candidates (top). A raw disease score is then calculated for each cell by quantifying the average expression of these candidate genes (middle). To assess statistical significance, matched control gene sets are generated to compute control scores, which are used to derive empirical p-value for each cell (bottom). This procedure enables robust identification of disease-associated cells.

The outputs of DPCGS include cell-specific TRSs, their associated p-values, and matched control scores. These results enable multiple downstream analyses. At the cell type level, DPCGS identifies which predefined cell populations are most strongly associated with the trait. At the gene level, it highlights trait-relevant genes specifically enriched in trait-associated subpopulations and links them to biological pathways. Finally, at the regulatory level, DPCGS predicts transcription factors that act as key regulators of trait-associated cellular states. Together, these analyses provide a comprehensive framework for dissecting the cellular and molecular mechanisms underlying complex traits and diseases at single-cell resolution.

### Performance evaluation using simulated data

To assess the efficacy and accuracy of DPCGS in identifying phenotype-associated cells, we first conducted performance evaluations using simulated datasets. For this purpose, we integrated real GWAS summary statistics with artificially constructed scRNA-seq datasets, focusing on the ability of DPCGS to distinguish trait-relevant cells.

As a benchmark trait, we selected monocyte count, a trait with strong heritability and biological relevance to immune system function. The GWAS summary statistics were derived from a large-scale study comprising 349,856 individuals, providing a robust foundation for analysis^13^. To construct the simulated scRNA-seq dataset, we generated single-cell profiles based on bulk expression data from fluorescence-activated cell-sorted hematopoietic cells (Methods). The dataset included 1,000 monocytes as positive examples (trait-associated cells) and 1,000 cells of unrelated types—T cells, B cells, dendritic cells (DCs), and natural killer (NK) cells—as negative examples (Fig. 2A). This design enabled a direct evaluation of DPCGS’s ability to discriminate target cells from non-target cells.

**Fig. 2.**
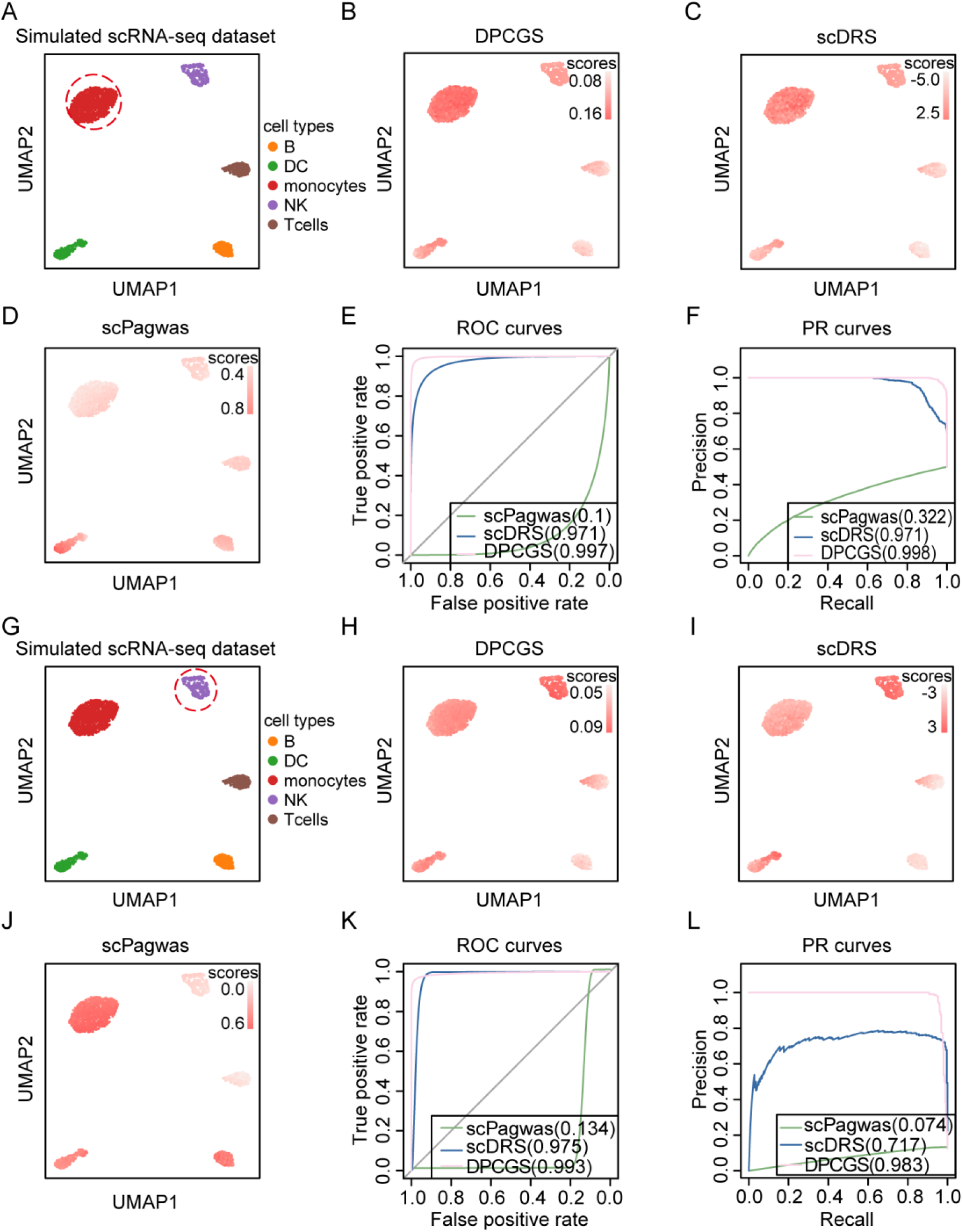
Evaluation of DPCGS in identifying trait-relevant cells in simulated data. (A) UMAP plot showing the cellular composition of the simulated scRNA-seq dataset for monocyte count; (B-D) Performance of DPCGS (B), scDRS (C), and scPagwas (D) in identifying monocyte count trait-associated cells; (E) Receiver operating characteristic (ROC) curves for the three methods in identifying monocyte count trait-associated cells; (F) Precision-recall (PR) curves for the three methods. (G) UMAP plot showing the cellular composition of the simulated scRNA-seq dataset for NK cell percentage; (H-J) Performance of DPCGS (H), scDRS (I), and scPagwas (J) in identifying NK cell percentage trait-associated cells; (K) Receiver operating characteristic (ROC) curves for the three methods in identifying NK cell percentage trait-associated cells; (L) Precision-recall (PR) curves for the three methods.

We then systematically compared DPCGS with existing cell-trait association methods, including scDRS and scPagwas. Prediction performance was quantified using the area under the receiver operating characteristic curve (AUROC) and the area under the precision-recall curve (AUPR). DPCGS consistently achieved superior results across both metrics (Fig. 2B-D). Specifically, DPCGS reached an AUROC of 0.997, outperforming scDRS (0.971) and scPagwas (0.100), and an AUPR of 0.998, markedly higher than scDRS (0.971) and scPagwas (0.322) (Fig. 2E, F). These findings demonstrate the strong discriminative power of DPCGS in identifying monocytes as trait-relevant cells.

To further evaluate the generalizability of DPCGS, we performed a second benchmark analysis on another immunologically relevant trait, NK cell percentage. The GWAS summary statistics, derived from 3,669 individuals, served as the genetic input^14^. The simulated dataset was constructed by defining NK cells as positive examples and other cell types (monocytes, T cells, B cells, and DCs) as negatives (Fig. 2G). Once again, DPCGS exhibited superior performance compared with scDRS and scPagwas. The AUROC of DPCGS was 0.993, higher than scDRS (0.975) and scPagwas (0.134), while the AUPR was 0.983, substantially exceeding scDRS (0.717) and scPagwas (0.074) (Fig. 2H-L).

Taken together, these results demonstrate that DPCGS consistently outperforms existing methods in distinguishing trait-associated from non-associated cells across different traits. By achieving high accuracy across diverse simulated scenarios, DPCGS shows strong potential as a versatile tool for identifying trait-relevant cellular subpopulations.

### Validation on real scRNA-seq datasets

To evaluate the performance of DPCGS in practical scenarios, we applied it to multiple representative scRNA-seq datasets. We first examined a bone marrow mononuclear cell (BMMC) scRNA-seq dataset that contains diverse immune cell types, aiming to identify cell populations associated with the monocyte count trait^15^ (Fig. 3A). Using DPCGS, scDRS, and scPagwas, we predicted and characterized trait-associated cells. Consistent with our simulation results, DPCGS demonstrated superior performance in distinguishing cells relevant to monocyte count (Fig. 3B-D). Specifically, DPCGS achieved an AUROC of 0.948, markedly higher than scDRS (0.921) and scPagwas (0.669) (Fig. 3E). Similarly, in terms of AUPR, DPCGS reached 0.825, substantially outperforming scDRS (0.685) and scPagwas (0.237) (Fig. 3F). These results confirm the strong accuracy and sensitivity of DPCGS in detecting trait-relevant cell populations from real data.

**Fig. 3.**
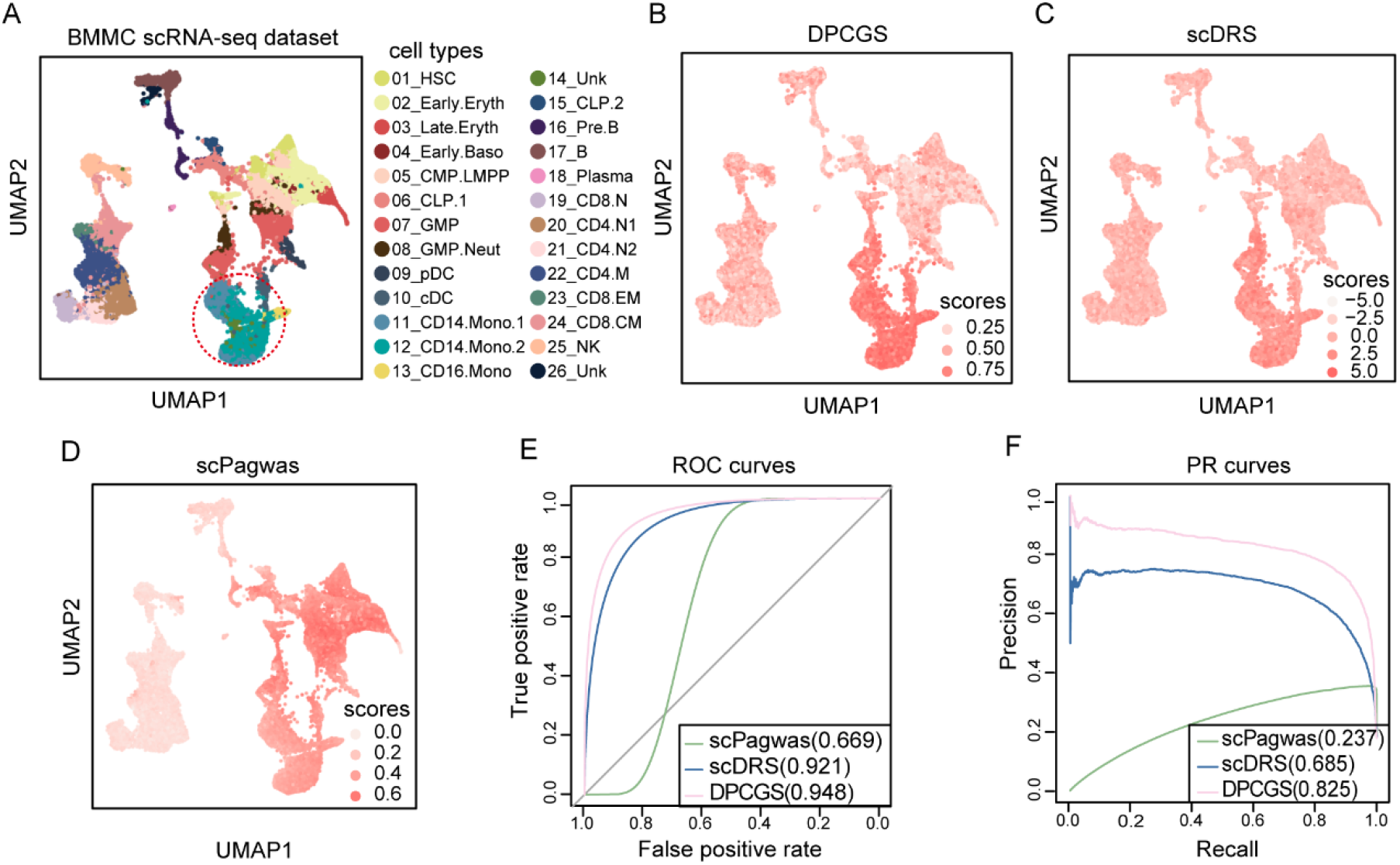
Evaluation of DPCGS in identifying monocyte count trait-relevant cells in BMMC data. (A) UMAP plot showing the cellular composition of the BMMC scRNA-seq dataset; (B-D) Performance of DPCGS (B), scDRS (C), and scPagwas (D) in identifying monocyte count trait-associated cells; (E) Receiver operating characteristic (ROC) curves for the three methods in identifying monocyte count trait-associated cells; (F) Precision-recall (PR) curves for the three methods in identifying monocyte count trait-associated cells.

To further assess the generalizability of DPCGS across different trait types, we investigated its performance on the NK cell percentage trait using a human peripheral blood immune cell scRNA-seq dataset^16^ (Fig. 4A). This dataset includes multiple immune cell types such as T cells, B cells, NK cells, and monocytes. DPCGS accurately identified cell groups associated with NK cell proportion and showed a clear advantage over competing methods (Fig. 4B-D. Quantitatively, DPCGS achieved an AUROC of 0.713, outperforming scDRS (0.681) and scPagwas (0.317) (Fig. 4E). In terms of AUPR, DPCGS also yielded the highest score compared to scDRS and scPagwas (Fig. 4F). These findings highlight the robustness of DPCGS in identifying NK cell-related traits and its applicability to different trait categories.

**Fig. 4.**
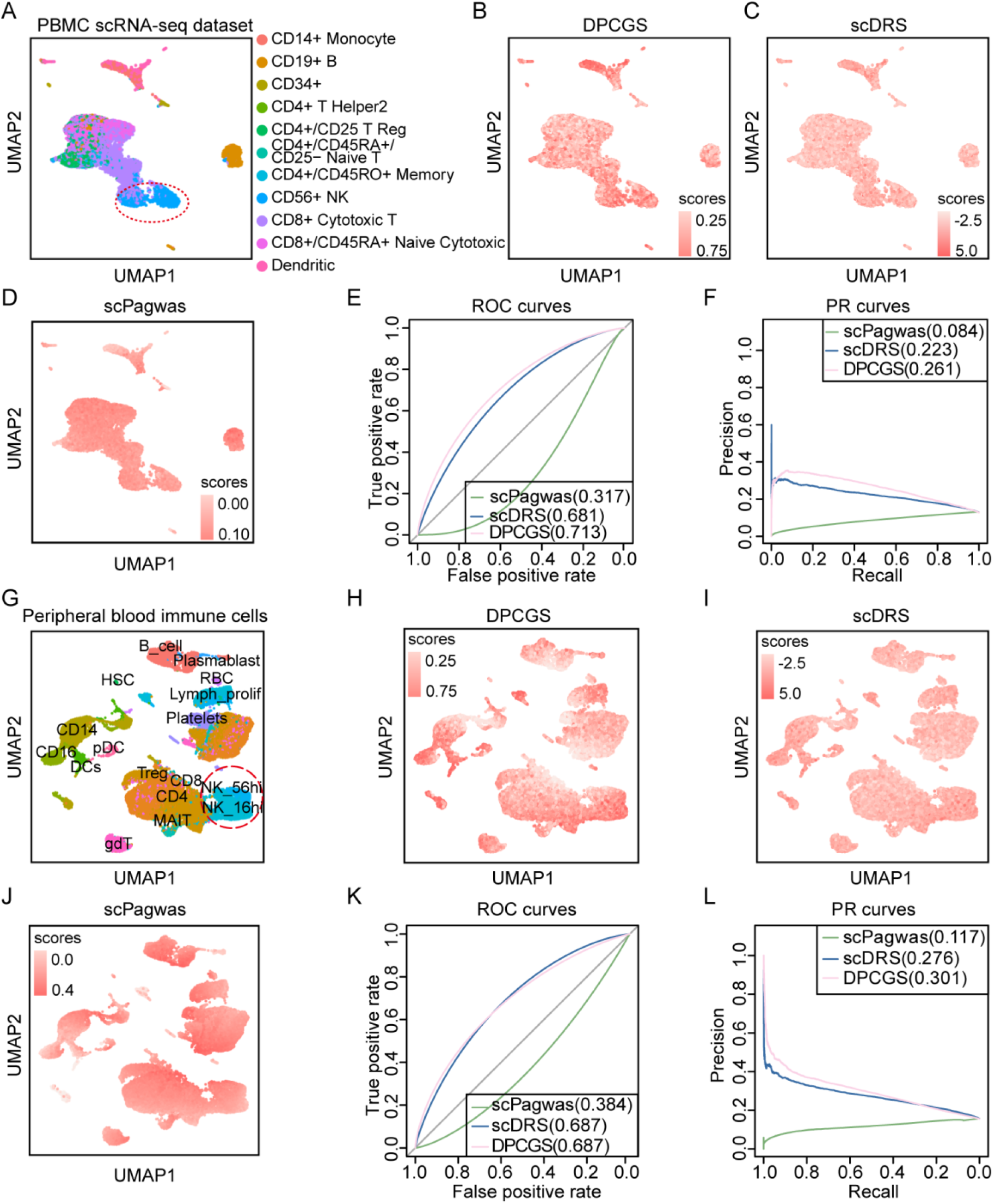
Evaluation of DPCGS in identifying NK cell percentage trait-relevant cells in PBMC and peripheral blood immune cell datasets. (A) UMAP plot showing the cellular composition of the PBMC scRNA-seq dataset; (B-D) Performance of DPCGS (B), scDRS (C), and scPagwas (D) in identifying NK cell percentage trait-associated cells; (E) Receiver operating characteristic (ROC) curves and (F) precision-recall (PR) curves comparing the three methods. (G) UMAP plot showing the cellular composition of the peripheral blood immune cell scRNA-seq dataset; (H-J) Identification of NK cell percentage trait-associated cells using DPCGS (H), scDRS (I), and scPagwas (J); (K) Receiver operating characteristic (ROC) curves and (L) precision-recall (PR) curves comparing the three methods.

We next evaluated the scalability of DPCGS using a large peripheral blood mononuclear cell (PBMC) scRNA-seq dataset containing 97,039 cells encompassing various immune lineages^17^ (Fig. 4G). The goal was to predict cells associated with the NK cell proportion trait and benchmark performance against scDRS and scPagwas. As shown in Fig. 4H-J, DPCGS maintained its superior performance. It achieved an AUROC of 0.687, comparable to scDRS (0.687) but substantially higher than scPagwas (0.384) (Fig. 4K). Importantly, its AUPR was also higher than that of both scDRS and scPagwas. (Fig. 4L). These results indicate that DPCGS remains accurate and reliable even in large-scale datasets, underscoring its scalability and robustness.

Together, these analyses demonstrate that DPCGS can robustly and accurately identify trait-associated cell populations across different real-world scRNA-seq datasets, regardless of dataset size or trait type. Compared to existing approaches, it shows superior accuracy, sensitivity, and robustness, highlighting its potential as a powerful computational framework for dissecting cellular heterogeneity and advancing immunological and disease-related research.

### Identification of heterogeneous cell subpopulations associated with AD

AD is a prevalent neurodegenerative disorder characterized by progressive neuronal loss and cognitive decline^18^. To better elucidate the cellular and molecular mechanisms underlying AD, we applied DPCGS by integrating single-nucleus RNA sequencing (snRNA-seq) data from the human entorhinal cortex with AD GWAS summary statistics^19, 20^. This analysis aimed to identify AD-associated cell subpopulations and their regulatory drivers.

The snRNA-seq dataset included six major cell types—neurons, astrocytes, oligodendrocytes, microglia, endothelial cells, and oligodendrocyte progenitor cells (OPCs) (Fig. 5A). By combining these transcriptomic profiles with GWAS data from 21,982 AD cases and 41,944 controls, we detected cell subpopulations with enriched genetic associations to AD (Fig. 5B). Among them, oligodendrocytes and astrocytes displayed the strongest enrichment signals (Fig. 5C). These findings are consistent with previous studies linking both cell types to neuroinflammation, amyloid plaque deposition, and neurodegeneration in AD^21, 22^. For example, Oligodendrocytes, crucial for myelination and axonal support, often show functional deficits in AD, while astrocytes, key regulators of neuronal homeostasis and synaptic plasticity, exhibit aberrant activation that may exacerbate neuronal damage and cognitive decline^23^.

**Fig. 5.**
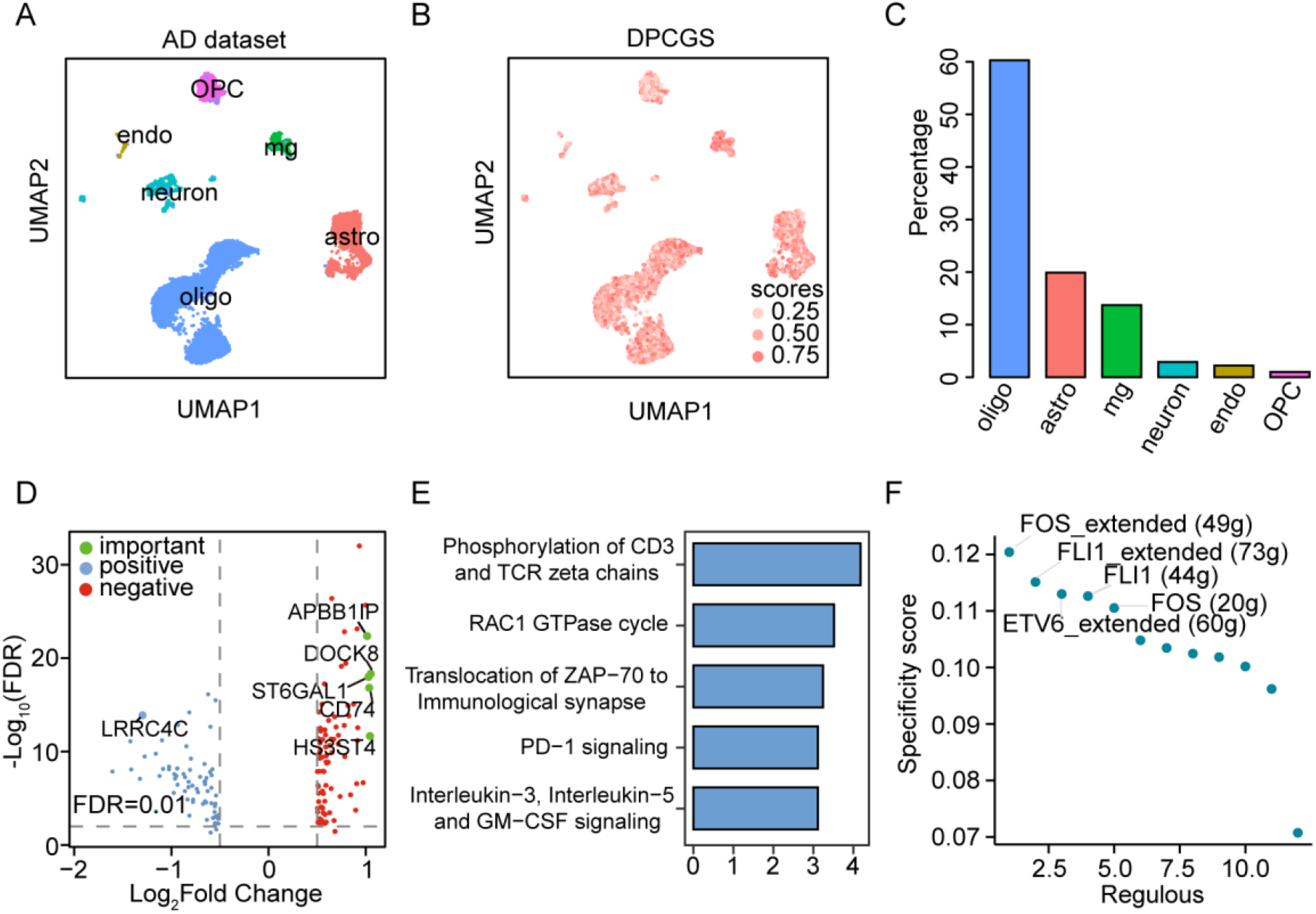
DPCGS identifies AD-associated cell subpopulations. (A) UMAP plot of human brain single-cell transcriptomic data, colored by cell type annotation. (B) UMAP plot showing AD phenotype relevance scores computed by DPCGS. (C) Cell type distribution of cells identified by DPCGS. (D) Differential expression analysis of genes between DPCGS-identified cells (labeled as 1) and other cells (labeled as 0) (absolute log2 fold change > 0.5, FDR < 0.01). (E) Top five Reactome pathways enriched in highly expressed genes in DPCGS-identified cells. (F) Ranking of regulators based on RSS in AD-associated cells identified by DPCGS.

Differential expression analysis of AD-associated subpopulations revealed markedly elevated expression of *CD74* (Fig. 5D and Supplementary Table S1). CD74, an immune regulatory molecule, has been implicated in several inflammatory diseases including AD^24, 25^. Its upregulation may contribute to heightened immune responses and neuroinflammation, suggesting a potential role as both a biomarker and a therapeutic target. Pathway enrichment analysis further demonstrated that the highly expressed genes were predominantly involved in immune-related processes such as T cell activation, signal transduction, and cytokine release (Fig. 5E and Supplementary Table S2). Dysregulation of these pathways is thought to drive persistent neuroinflammation, thereby accelerating neuronal injury and AD progression^26, 27^.

Transcriptional regulatory analysis highlighted FOS and FLI1 as key transcription factors within the AD-associated subpopulations (Fig. 5F and Supplementary Table S3). FOS, a member of the AP-1 family, is widely implicated in stress responses, immune regulation, and synaptic function, with abnormal expression linked to neuronal injury and amyloid pathology in AD. FLI1, an ETS family factor, is primarily known for roles in hematopoiesis, angiogenesis, and immune function. Although less studied in AD, its dysregulation may contribute to blood-brain barrier dysfunction and enhanced immune activation, thereby aggravating neuroinflammation.

Together, these findings demonstrate that DPCGS effectively identifies AD-associated cell subpopulations and their molecular regulators. Our results highlight the involvement of oligodendrocytes and astrocytes in AD and suggest CD74, FOS, and FLI1 as potential mediators of disease pathology.

### Detection of cells related to asthma

Asthma is a common chronic respiratory disease affecting more than 300 million people worldwide, characterized by persistent airway inflammation, hyperresponsiveness, and remodeling^28^. Its etiology is multifactorial, involving both genetic and environmental factors. To investigate the cellular basis of asthma, we applied the DPCGS framework by integrating single-cell transcriptomic profiles from human lung tissues with large-scale GWAS summary statistics^29^ (Fig. 6A). This analysis enabled systematic identification of asthma-associated cell subpopulations and exploration of their potential roles in pathogenesis.

**Fig. 6.**
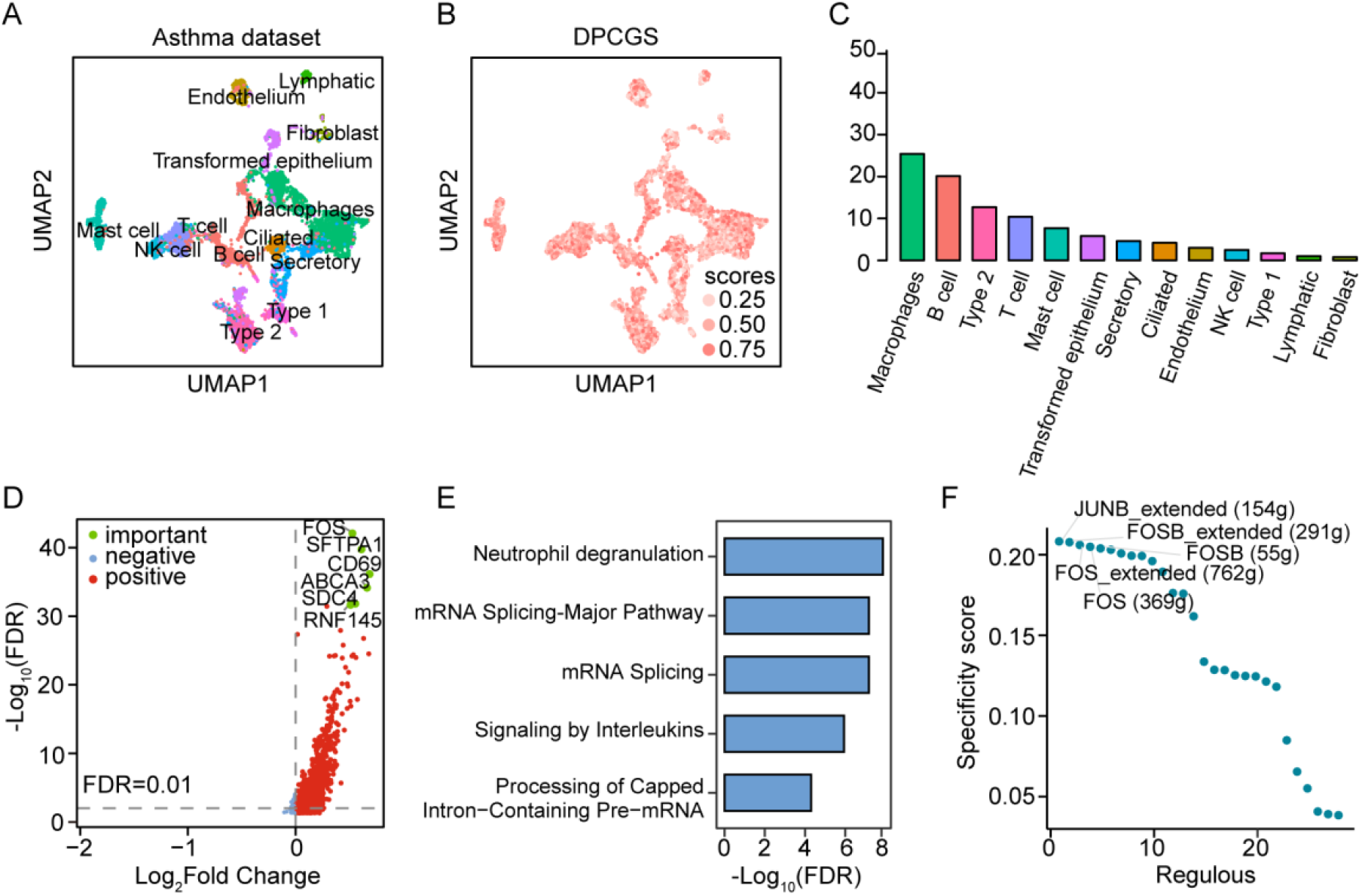
DPCGS identifies asthma-associated cell subpopulations. (A) UMAP plot of human lung single-cell transcriptomic data, colored by cell type annotation. (B) UMAP plot showing asthma phenotype relevance scores computed by DPCGS. (C) Cell type distribution of DPCGS-identified cells. (D) Differential expression analysis of genes between DPCGS-identified cells (labeled as 1) and other cells (labeled as 0) (absolute log2 fold change > 0, FDR < 0.01). (E) Top five Reactome pathways enriched in highly expressed genes in DPCGS-identified cells. (F) Ranking of regulators based on RSS in asthma-associated cells identified by DPCGS.

We first examined scRNA-seq data from lung tissues of asthma patients and healthy controls, encompassing 13 annotated cell types including lymphocytes, endothelial cells, fibroblasts, epithelial cells, macrophages, and ciliated cells (Fig. 6A). DPCGS identified significantly elevated asthma-association scores in macrophages and B cells (Fig. 6B, C). This observation is consistent with previous studies implicating both cell types in chronic airway inflammation and immune dysregulation during asthma development^30, 31^.

Differential gene expression analysis further revealed distinct molecular signatures within asthma-associated subpopulations (Fig. 6D and Supplementary Table S4). Among these, *FOS* emerged as the most significantly upregulated gene. As a key component of the AP-1 transcription factor family, FOS regulates cell proliferation, stress responses, and differentiation^32^. In the context of asthma, FOS has been implicated in airway remodeling by promoting smooth muscle proliferation, collagen deposition, and epithelial changes^33, 34^. These findings highlight FOS as a potential molecular driver of disease progression and a candidate therapeutic target.

Pathway enrichment analysis showed that highly expressed genes in asthma-associated cell subsets were significantly enriched in pathways related to neutrophil degranulation (Fig. 6E and Supplementary Table S5). Neutrophils are known to play critical roles in severe or exacerbated asthma, where they release antimicrobial proteins, cytokines, and inflammatory mediators (e.g., myeloperoxidase, elastase, IL-8) that amplify airway inflammation and hyperreactivity^35, 36^. These results suggest that targeting neutrophil accumulation or inhibiting degranulation processes may represent promising therapeutic strategies.

To further dissect the regulatory landscape, we performed transcriptional regulatory analysis on asthma-associated subpopulations (Fig. 6F and Supplementary Table S6). We identified AP-1 family transcription factors—JUNB, FOS, and FOSB—as key regulators with strong cell-type specificity^37^. These factors are known mediators of immune activation, stress responses, and airway function^38^. Their upregulation in asthma-associated subpopulations likely contributes to sustained immune activation and airway remodeling, consistent with the differential expression results.

In summary, our integrative analysis using DPCGS identified macrophages and B cells as key immune subpopulations associated with asthma. The regulatory role of AP-1 transcription factors provides further insight into disease mechanisms, offering potential molecular targets for therapeutic intervention. Together, these findings illustrate the power of DPCGS in delineating cell type-specific mechanisms underlying complex respiratory diseases.

## Discussion

In this study, we introduced DPCGS, a computational framework that integrates single-cell transcriptomic profiles with GWAS summary statistics to identify trait-associated cell subpopulations and their molecular regulators. Through systematic evaluations on both simulated and real datasets, as well as applications to complex diseases including AD and asthma, we demonstrated that DPCGS consistently outperforms existing approaches in accuracy, sensitivity, and robustness.

Our benchmarking analyses using simulated data established the strong discriminative power of DPCGS in distinguishing trait-relevant from non-relevant cells. Compared to scDRS and scPagwas, DPCGS showed superior performance across multiple traits, highlighting its reliability across diverse contexts. Importantly, validation on real scRNA-seq datasets further confirmed its robustness and scalability. Even in large-scale datasets containing nearly 100,000 cells, DPCGS maintained high predictive accuracy, demonstrating its applicability to increasingly large single-cell studies.

Beyond quantitative evaluation, the application of DPCGS to disease-relevant datasets provided new insights into disease pathogenesis. In AD, we identified oligodendrocytes and astrocytes as key disease-associated cell populations, consistent with prior studies, while also highlighting candidate regulators such as CD74, FOS, and FLI1 that may contribute to immune dysregulation and neuroinflammation. Similarly, in asthma, DPCGS pinpointed macrophages and B cells as critical immune subsets, with transcriptional regulation by AP-1 family members (e.g., JUNB and FOSB). These findings not only corroborate known mechanisms but also uncover novel regulatory signatures, suggesting potential biomarkers and therapeutic targets. Collectively, these applications demonstrate the effectiveness of DPCGS in linking GWAS findings to functional insights specific to cell types.

Despite these promising results, several limitations should be acknowledged. First, the accuracy of DPCGS depends on the quality of GWAS summary statistics and the mapping of SNPs to genes, which may not fully capture long-range regulatory interactions. Second, scRNA-seq data remain subject to technical noise and batch effects, which could affect disease score estimation. Incorporating complementary data modalities, such as chromatin accessibility (scATAC-seq) or spatial transcriptomics, may further enhance the resolution and interpretability of DPCGS analyses. Finally, while our applications highlight associations between cell subpopulations and disease, causal inference requires integration with perturbation data or experimental validation.

In conclusion, DPCGS provides a powerful and versatile framework for dissecting the cellular and molecular basis of complex traits and diseases at single-cell resolution. By linking genetic variation to cell-type-specific expression programs and regulatory networks, it enables the systematic discovery of disease-relevant cellular contexts and candidate drivers. With continued refinement and integration of multi-omics data,

DPCGS holds great promise for advancing precision medicine by identifying novel biomarkers and therapeutic targets.

## Methods

### Description of DPCGS

The DPCGS algorithm is designed to quantify the association between individual cells and disease-relevant molecular signals using GWAS data. The algorithm consists of three major steps: (1) construction of a disease-associated gene set based on GWAS summary statistics; (2) calculation of disease scores and background control scores for each cell; and (3) evaluation of statistical significance by comparing normalized disease scores with the empirical distribution of control scores. Each of these steps is detailed below.

### Construction of Disease-Associated Gene Set

We employed MAGMA framework to perform gene-level association analysis using GWAS summary statistics for the trait or disease of interest^39^. For a given gene, MAGMA evaluates its statistical significance by calculating the test statistic:

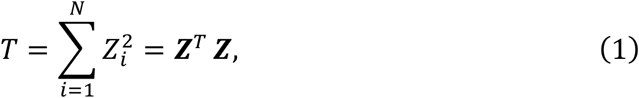

where *N* denotes the number of SNPs mapped to a gene, including those located within the gene body and ±10 kb flanking regions, and *Z*_*i*_ is the standard normal deviate corresponding to the marginal p-value of SNP _*i*_ computed as *Z*_*i*_ = ϕ^−1^(*p*_*i*_), where ϕ is the standard normal cumulative distribution function.

MAGMA assumes a multivariate normal distribution ***Z***∼*MVN*(0, ***S***), where ***S*** is the Linkage disequilibrium (LD) matrix among SNPs. The LD matrix was calculated using European population data from the 1,000 Genomes Project Phase 3. It can be decomposed as ***S*** = ***QAQ***^*T*^, where ***A*** = *d*_*i*_*ag*(λ_1_, λ_1_, …, λ_*N*_) contains the eigenvalues λ_*l*_, and ***Q*** is the orthogonal matrix of eigenvectors. Using the transformation ***D*** = ***A***^−0.5^***Q***^*T*^***Z***∼*MVN*(0, ***I***_*k*_), the test statistic becomes:

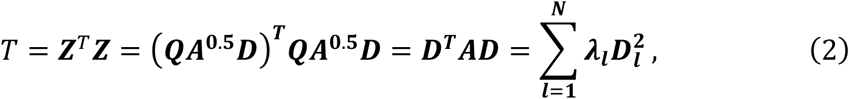

where *D*_*i*_∼*N*(0,1), and thus *T* follows a linear combination of independent 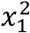 distributions. Genes are ranked based on their *T*-values to produce a MAGMA output gene list *G*_*MAGMA*_.

Let ***E*** ∈ ℝ^*m*×*n*^ denote the gene expression matrix with *m* genes and *n* cells. We intersect the MAGMA gene list with the gene expression matrix to obtain the filtered gene set:

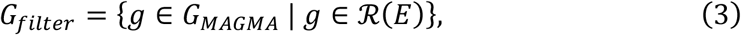

where ℛ(***E***) denotes the set of gene names in the expression matrix. The final disease-associated gene set *G* = {*g*_1_, *g*_2_, …, *g*_1000_} consists of the top 1,000 genes from *Gfilter* .

### Calculation of Disease Scores

For each cell *j* ∈ {1,2, …, *n*}, we calculated a disease score based on the mean expression of the genes in the disease-associated gene set *G*, and standardized it against multiple control gene sets as follows:

### Compute target gene set score

The raw disease score is defined as:

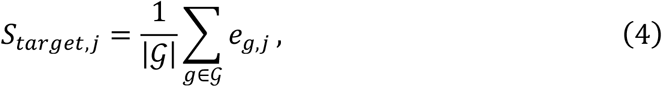

where |*G*| is the number of matched genes in the expression matrix and *e*_*g,j*_ representing the expression of gene *g* in cell *j*.

### Compute background gene set scores

We randomly sampled *M* background gene sets *B*^(*m*)^ of the same size as *B*^(*m*)^, and computed their mean expression per cell:

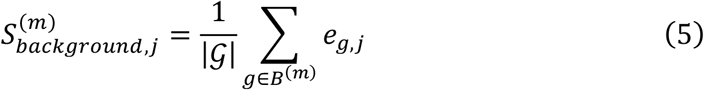

### Calculate standardized target gene set scores

To normalize the target score, we subtracted the average background score:

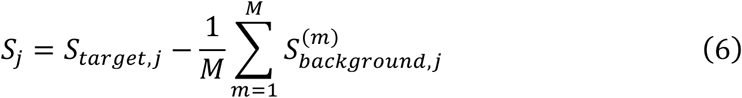

### Transform standardized scores to a sigmoid-based range

We normalized *S*_*j*_ across all cells to have zero mean and unit variance:

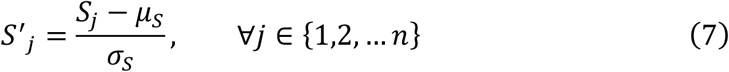

where 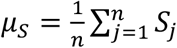 and 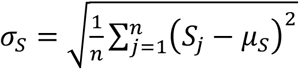

Finally, a sigmoid transformation was applied to map normalized scores to the range (0,1):

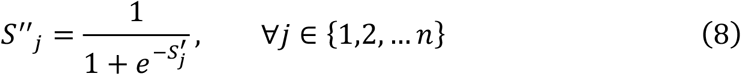

The transformed score vector ***S***^′′^ = (*S*^′′^_1_, *S*^′′^_2_, ⋯ *S*^′′^_*n*_) represents the final disease module scores per cell.

### Statistical Significance and Multiple Testing Correction

To evaluate the statistical significance of each cell’s disease association, we generated 1,000 random control gene sets by sampling 1,000 genes from the expression matrix *E* uniformly at random:

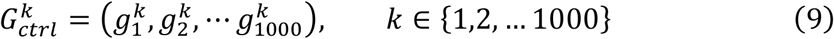

Each control gene set was used to compute a corresponding control score 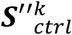 using the same method as the disease score. For each cell *j*, we estimated an empirical p-value by comparing its disease score *S*^′′^_*j*_ to the empirical distribution of control scores across the 1,000 permutations:

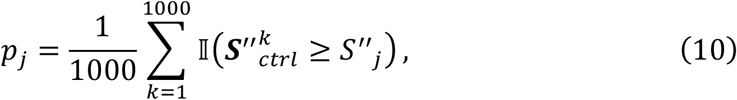

where I(.) is the indicator function. Finally, Benjamini-Hochberg correction was applied to the set of p-values 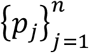 to control the False Discovery Rate (FDR).

### Simulations

To evaluate the performance of DPCGS in identifying individual cells associated with traits such as monocyte count or NK cell percentage, we simulated a scRNA-seq dataset containing five distinct cell types using the scDesign2 package^40^. The simulated cell types included monocytes, dendritic cells (DCs), B cells, NK cells, and T cells. Among these, DCs—differentiated from monocytes—were selected as a cell type unrelated to the monocyte-associated trait, serving as a potential confounding factor for distinguishing monocytes from other simulated cells.

In the model fitting stage, we used fluorescence-activated cell sorted (FACS) blood system samples (GEO accession: GSE107011) as the reference dataset to fit a multivariate generative model. Since the reference dataset comprised five sorted cell types, we partitioned the data into five subsets based on cell type and fitted a cell type-specific model for each subset.

In the data generation stage, synthetic scRNA-seq data were sampled from the fitted models to represent both phenotype-associated cell populations (e.g., monocytes) and phenotype-unrelated cell populations (e.g., DCs, B cells, NK cells, and T cells). The final synthetic dataset contained 2,000 cells, with the following proportions: monocytes (0.50), DCs (0.125), B cells (0.125), NK cells (0.125), and T cells (0.125).

### scRNA-seq data and pre-processing

We analyzed five scRNA-seq or snRNA-seq datasets. For blood cell-related traits, we collected a human bone marrow mononuclear cell (BMMC) scRNA-seq dataset (n=35,582) ^15^, a human peripheral blood mononuclear cell (PBMC) scRNA-seq dataset (n=97,039), and a human peripheral blood immune cell scRNA-seq dataset (n=32,738)^16, 17^. For AD-related analysis, we used a human brain snRNA-seq dataset (n=13,214) ^19^. Additionally, we included an scRNA-seq dataset of freshly resected human lung tissue (n=10,360) ^29^.

For all scRNA-seq datasets, we first removed low-quality cells with a mitochondrial gene count fraction greater than or equal to 0.2, as these cells were likely undergoing apoptosis or lysis. Gene expression counts were normalized to a total of 10,000 counts per cell, followed by log-transformation. The resulting normalized gene expression matrices were further standardized by centering each gene’s expression values (mean subtraction) and scaling to unit variance, thereby removing scale effects and enhancing comparability across genes. This standardization step facilitates downstream analyses such as dimensionality reduction and clustering.

Highly variable genes (HVGs) were identified to capture the most biologically informative features, and the top 3,000 HVGs were selected for subsequent analysis. Principal component analysis (PCA) was then applied to the standardized expression matrix, and the top 50 principal components were retained to summarize the major axes of variation in the data. To visualize the cellular heterogeneity, we performed Uniform Manifold Approximation and Projection (UMAP) on the PCA-reduced data. A k-nearest neighbor (kNN) graph was constructed based on the principal components to assess cell-to-cell similarity, and graph-based clustering was performed with a resolution parameter of 0.3 to identify distinct cell populations.

### GWAS summary statistics

All GWAS summary statistics were obtained from publicly available projects hosted by the IEU Open GWAS database (https://gwas.mrcieu.ac.uk/). These datasets comprised two hematological traits, namely monocyte count (ukb-d-30130_irnt)^13^ and NK cell percentage (ebi-a-GCST90001647)^14^, together with AD (ieu-b-2)^20^ and asthma (ukb-d-J10_ASTHMA).

Based on previous evidence that most expression quantitative trait loci (eQTLs) are located within a 20 kb window centered on the transcription start site of genes^7^, we adopted a default ±20 kb window in the DPCGS algorithm to assign SNPs from the GWAS summary statistics to their corresponding genes. In subsequent analyses, LD between available SNPs was calculated using the European-ancestry reference panel from Phase 3 of the 1000 Genomes Project^41^. Because the major histocompatibility complex region (Chr6: 25-35 Mbp) exhibits extensive LD, this region was excluded from all analyses to avoid confounding effects^42^.

### Comparison of DPCGS with existing approaches

To evaluate the effectiveness of the proposed DPCGS method, we compared it with several alternative approaches for identifying disease-associated cell subpopulations. A brief description of these comparative methods is provided below.

The scDRS method links individual cells to diseases by combining scRNA-seq gene expression data with GWAS-derived polygenic risk, evaluating whether putative disease-associated genes exhibit elevated expression in a cell compared to genes with comparable overall expression patterns across all cells^8^. scPagwas is a pathway-based polygenic regression method designed to integrate scRNA-seq data with GWAS results to identify cellular contexts associated with complex diseases and traits^9^. The method employs a linear regression framework to relate GWAS signals to pathway activation profiles derived from scRNA-seq data, thereby identifying trait-associated gene sets and inferring the most relevant cell subpopulations.

A detailed performance comparison between these methods and DPCGS, focusing on their ability to identify disease-associated cell subpopulations, is presented in the Results section.

### Differential expression and pathway enrichment analysis

Differentially expressed genes (DEGs) were identified using the Wilcoxon rank-sum test, implemented via the FindMarkers function in the Seurat R package^43^. Genes were considered significantly differentially expressed if they met both a minimum absolute log_2_ fold change (|log_2_FC|) threshold of 0.5 and a maximum Bonferroni-adjusted p-value of 0.05. The resulting DEG set was subsequently subjected to pathway enrichment analysis using the Reactome database. This analysis was performed with the ReactomePA R package^44^, applying the hypergeometric test to evaluate the statistical significance of each pathway and thereby identify potential biological processes and signaling pathways associated with the DEGs.

### Analysis of gene regulatory networks

Gene regulatory network analysis was conducted using the SCENIC framework^45^ to explore the regulatory architecture of the cell subpopulations identified by the DPCGS method. First, co-expression modules linking transcription factors (TFs) to their potential target genes were identified from the single-cell transcriptomic data. These modules were then refined using the RcisTarget R package, which infers direct TF-target interactions by identifying TF-binding motifs significantly enriched in the candidate target genes based on genome-wide motif-ranking databases. For each regulon, defined as a TF and its direct target genes, regulon activity scores were computed to quantify its activity across individual cells. Finally, an entropy-based approach was applied to calculate the Regulon Specificity Score (RSS) for each regulon within each subpopulation, enabling the identification of key regulons with high specificity in DPCGS-defined cell clusters.

## Supporting information

Description of Supplementary Files

## Data availability

All GWAS summary statistics used in this study were obtained from the publicly available IEU OpenGWAS Project. Specifically, summary statistics for monocyte count were accessed at https://opengwas.io/datasets/ukb-d-30130_irnt, and those for NK cell percentage at https://opengwas.io/datasets/ebi-a-GCST90001647. Summary statistics for AD were obtained from https://gwas.mrcieu.ac.uk/datasets/ieu-b-2/, and those for asthma from https://gwas.mrcieu.ac.uk/datasets/ukb-d-J10_ASTHMA/. The bone marrow mononuclear cell (BMMC) single-cell RNA-seq dataset was retrieved from https://jeffgranja.s3.amazonaws.com/MPAL-10x/Supplementary_Data/Healthy-Data/scRNA-Healthy-Hematopoiesis-191120.rds, and the peripheral blood mononuclear cell (PBMC) dataset was obtained from the 10x Genomics GitHub repository (https://github.com/10XGenomics/single-cell-3prime-paper/tree/master/pbmc68k_analysis). The peripheral blood immune cell scRNA-seq dataset was downloaded from ArrayExpress (accession: E-MTAB-10026). Human brain single-nucleus RNA-seq data were obtained from GEO under accession GSE138852, and single-cell transcriptomic data from human lung tissue were downloaded from GEO under accession GSE130148.

## Code availability

The code to reproduce the DPCGS method and all results of the paper is available on GitHub (https://github.com/Chonghui-Liu/DPCGS).

## Acknowledgements

We gratefully acknowledge the computational resources provided by Kunming University of Science and Technology, which were essential for the large-scale benchmarking and application of our DPCGS framework. We also thank the researchers and consortia who generated and shared the public single-cell and GWAS datasets that formed the foundation for our methodological development and biological discoveries.

## Author contributions

C.L. conceived the overall research framework. The algorithm was developed and applied by C.L. Interpretation of the results was a joint effort by B.Y., B.S., J.L., R.Z., P.Y., B.W., Y.X., S.Y., N.Y., L.M., and Q.L. The study was supervised by S.D. and Y.Z. The manuscript was prepared by C.L., S.D. and Y.Z., with input and critical revisions provided by all co-authors. All authors have read and approved the final version of the manuscript.

## Supporting Information

Supporting Information is available in the attachment.

## References

1. Timpson, N. J., Greenwood, C. M. T., Soranzo, N., Lawson, D. J. & Richards, J. B. Genetic architecture: the shape of the genetic contribution to human traits and disease. Nat. Rev. Genet. 19, 110–124 (2018).

2. Spielmann, M. & Mundlos, S. Looking beyond the genes: the role of non-coding variants in human disease. Hum. Mol. Genet. 25, R157–R165 (2016).

3. Trubetskoy, V., et al. Mapping genomic loci implicates genes and synaptic biology in schizophrenia. Nature 604, 502–508 (2022).

4. Locke, A. E., et al. Genetic studies of body mass index yield new insights for obesity biology. Nature 518, 197–206 (2015).

5. Pers, T. H., et al. Biological interpretation of genome-wide association studies using predicted gene functions. Nat. Commun. 6, 5890 (2015).

6. Papalexi, E. & Satija, R. Single-cell RNA sequencing to explore immune cell heterogeneity. Nat. Rev. Immunol. 18, 35–45 (2018).

7. Hekselman, I. & Yeger-Lotem, E. Mechanisms of tissue and cell-type specificity in heritable traits and diseases. Nat. Rev. Genet. 21, 137–150 (2020).

8. Zhang, M. J., et al. Polygenic enrichment distinguishes disease associations of individual cells in single-cell RNA-seq data. Nat. Genet. 54, 1572–1580 (2022).

9. Ma, Y., et al. Polygenic regression uncovers trait-relevant cellular contexts through pathway activation transformation of single-cell RNA sequencing data. Cell Genomics 3, (2023).

10. Finucane, H. K., et al. Heritability enrichment of specifically expressed genes identifies disease-relevant tissues and cell types. Nat. Genet. 50, 621–629 (2018).

11. Calderon, D., et al. Inferring relevant cell types for complex traits by using single-cell gene expression. Am. J. Hum. Genet. 101, 686–699 (2017).

12. Watanabe, K., Umićević Mirkov, M., de Leeuw, C. A., van den Heuvel, M. P. & Posthuma, D. Genetic mapping of cell type specificity for complex traits. Nat. Commun. 10, 3222 (2019).

13. Elsworth, B., et al. The MRC IEU OpenGWAS data infrastructure. BioRxiv, 2020–2008 (2020).

14. Orrù, V., et al. Complex genetic signatures in immune cells underlie autoimmunity and inform therapy. Nat. Genet. 52, 1036–1045 (2020).

15. Granja, J. M., et al. Single-cell multiomic analysis identifies regulatory programs in mixed-phenotype acute leukemia. Nat. Biotechnol. 37, 1458–1465 (2019).

16. Stephenson, E., et al. Single-cell multi-omics analysis of the immune response in COVID-19. Nat. Med. 27, 904–916 (2021).

17. Zheng, G. X. Y., et al. Massively parallel digital transcriptional profiling of single cells. Nat. Commun. 8, 14049 (2017).

18. Jones, D., et al. A computational model of neurodegeneration in Alzheimer’s disease. Nat. Commun. 13, 1643 (2022).

19. Grubman, A., et al. A single-cell atlas of entorhinal cortex from individuals with Alzheimer’s disease reveals cell-type-specific gene expression regulation. Nat. Neurosci. 22, 2087–2097 (2019).

20. Kunkle, B. W., et al. Genetic meta-analysis of diagnosed Alzheimer’s disease identifies new risk loci and implicates Aβ, tau, immunity and lipid processing. Nature genetics 51, 414–430 (2019).

21. Sadick, J. S., et al. Astrocytes and oligodendrocytes undergo subtype-specific transcriptional changes in Alzheimer’s disease. Neuron 110, 1788–1805 (2022).

22. Park, H., et al. Single-cell RNA-sequencing identifies disease-associated oligodendrocytes in male APP NL-G-F and 5XFAD mice. Nat. Commun. 14, 802 (2023).

23. Simons, M. & Nave, K.-A. Oligodendrocytes: myelination and axonal support. Cold Spring Harbor Perspect. Biol. 8, a020479 (2016).

24. Su, H. T., Na, N., Zhang, X. D. & Zhao, Y. The biological function and significance of CD74 in immune diseases. Inflammation Res. 66, 209–216 (2017).

25. Bryan, K. J., et al. Expression of CD74 is increased in neurofibrillary tangles in Alzheimer’s disease. Mol. Neurodegener. 3, 13 (2008).

26. Bettcher, B. M., Tansey, M. G., Dorothée, G. & Heneka, M. T. Peripheral and central immune system crosstalk in Alzheimer disease — a research prospectus. Nat. Rev. Neurology 17, 689–701 (2021).

27. Wu, K. M., et al. The role of the immune system in Alzheimer’s disease. Ageing Res. Rev. 70, 101409 (2021).

28. Boonpiyathad, T., Sözener, Z. C., Satitsuksanoa, P. & Akdis, C. A. Immunologic mechanisms in asthma. Semin. Immunol. 46, 101333 (2019).

29. Vieira Braga, F. A., et al. A cellular census of human lungs identifies novel cell states in health and in asthma. Nat. Med. 25, 1153–1163 (2019).

30. van der Veen, T. A., de Groot, L. E. S. & Melgert, B. N. The different faces of the macrophage in asthma. Curr. Opin. Pulm. Med. 26, 62–68 (2020).

31. Alturaiki, W. The roles of B cell activation factor (BAFF) and a proliferation-inducing ligand (APRIL) in allergic asthma. Immunol. Lett. 225, 25–30 (2020).

32. Ordway, J. M., Fenster, S. D., Ruan, H. & Curran, T. A transcriptome map of cellular transformation by the fos oncogene. Mol. Cancer 4, 19 (2005).

33. Varricchi, G., et al. Biologics and airway remodeling in severe asthma. Allergy77, 3538–3552 (2022).

34. He, Y. Y., et al. The Fra-1: Novel role in regulating extensive immune cell states and affecting inflammatory diseases. Front. Immunol. 13, 954744 (2022).

35. Hammad, H. & Lambrecht, B. N. The basic immunology of asthma. Cell 184, 1469–1485 (2021).

36. Li, Y. M., Yang, T. Y. & Jiang, B. H. Neutrophil and neutrophil extracellular trap involvement in neutrophilic asthma: A review. Medicine 103, e39342 (2024).

37. Wagner, E. F. Bone development and inflammatory disease is regulated by AP-1 (Fos/Jun). Ann. Rheum. Dis. 69, i86–i88 (2010).

38. Hartenstein, B., et al. Th2 cell-specific cytokine expression and allergen-induced airway inflammation depend on JunB. The EMBO journal (2002).

39. De Leeuw, C. A., Mooij, J. M., Heskes, T. & Posthuma, D. MAGMA: generalized gene-set analysis of GWAS data. PLoS Comput. Biol. 11, e1004219 (2015).

40. Sun, T., Song, D., Li, W. V. & Li, J. J. scDesign2: a transparent simulator that generates high-fidelity single-cell gene expression count data with gene correlations captured. Genome Biol. 22 (2021).

41. Altshuler, D. M., et al. A global reference for human genetic variation. Nature 526, 68–74 (2015).

42. Kilpinen, H., et al. Common genetic variation drives molecular heterogeneity in human iPSCs. Nature 546, 370–375 (2017).

43. Stuart, T., et al. Comprehensive Integration of Single-Cell Data. Cell 177, 1888–1902.e1821 (2019).

44. Yu, G. & He, Q.-Y. ReactomePA: an R/Bioconductor package for reactome pathway analysis and visualization. Mol. BioSyst. 12, 477–479 (2016).

45. Aibar, S., et al. SCENIC: single-cell regulatory network inference and clustering. Nat. Methods 14, 1083–1086 (2017).

