## Supplementary material for "DPCGS: a computational framework for linking GWAS to single-cell transcriptomics in complex traits and diseases": Description of Supplementary Files

File Name: Supplementary Table S1

Description: Differentially expressed genes between DPCGS-identified cells associated with AD versus all other cells

File Name: Supplementary Table S2

Description: Reactome pathways enrichment of upregulated genes in DPCGS-identified cells associated with AD

File Name: Supplementary Table S3

Description: The specificity scores of regulons in DPCGS-identified cells associated with AD

File Name: Supplementary Table S4

Description: Differentially expressed genes between DPCGS-identified cells associated with asthma versus all other cells

File Name: Supplementary Table S5

Description: Reactome pathways enrichment of upregulated genes in DPCGS-identified cells associated with asthma

File Name: Supplementary Table S6

Description: The specificity scores of regulons in DPCGS-identified cells associated with asthma
